# Designing spatial capture–recapture surveys for multiple populations

**DOI:** 10.64898/2026.09.22.753485

**Authors:** Greg Distiller, Ian Durbach

## Abstract

Recent work on the design of spatial capture-recapture (SCR) studies has largely focused on settings with a single set of movement and detection parameters. In many applied contexts, however, populations are naturally stratified—for example, when different sexes exhibit markedly different movement scales and encounter processes, or when more than one focal species is monitored. Designs that perform well for one group may be inefficient or suboptimal for others, despite the substantial logistical effort typically required to implement an SCR survey.

We describe several potential SCR design strategies for surveying multiple groups, and compare these using simulation. Designs considered include regular and optimised grids, clustered and lacework designs, and designs generated using genetic algorithms, including new approaches that optimise information from both groups and a recently published two-stage design. Using proximity detectors such as camera traps, we evaluate design performance in terms of precision, bias, and coverage, and examine how performance changes with the number of available resources.

Our results show that multi-group SCR surveys can perform well even when focal groups differ greatly in movement scale and encounter rate, provided that some simple guidelines are followed. Systematic layouts accommodating more than one characteristic spacing e.g. cluster designs, performed consistently well and are a reliable default option. Regular grid designs with a single spacing between detectors are feasible when differences between groups are moderate but performed poorly as differences increased. Establishing when these designs become problematic adds unnecessary risk to the design process. Algorithmic designs did not outperform systematic designs even in highly-skewed scenarios expected to favour them. In practice, when there is more than one focal group we recommend identifying the smallest and largest plausible movement scales and rejecting potentially pathological designs, primarily those that cannot generate recaptures at the smaller scale or cover a substantial area at the larger scale.

## 1 Introduction

Spatial capture–recapture (SCR) models have become the standard framework for modelling animal density. By incorporating information on where individuals are detected, SCR models allow population density to be estimated directly (Efford 2004; Borchers and Efford 2008). Despite the central role of detector placement in determining the quality of SCR data, comparatively little attention has been devoted to study design. Early guidance consisted largely of heuristic rules, often inherited from non-spatial capture–recapture literature. However one of the key insights from early SCR design studies was that SCR models are substantially more robust to variation in trap-array configuration and extent than traditional non-spatial capture–recapture models because they explicitly model the spatial relationship between detections and individual activity centres (Sollmann et al. 2012; Sun et al. 2014).

Simulation and empirical studies have shown that SCR density estimates are generally robust to variation in detector spacing and array geometry, remaining approximately unbiased across a wide range of designs provided that the detector array is sufficiently large relative to the scale of animal movement (Sollmann et al. 2012; Tobler and Powell 2013; Sun et al. 2014; Fleming et al. 2021). These studies also highlight the importance of obtaining adequate numbers of spatial recaptures to support reliable estimation of movement parameters and hence density. Although SCR estimators appear robust with respect to bias, the precision of density estimates is influenced by the number of spatial recaptures and the information available for estimating animal movement (Palmero et al. 2023). A review of applied SCR studies found that survey design was seldom evaluated explicitly and that imprecise density estimates were common (Green et al. 2020), highlighting the need for more rigorous survey design evaluation. Together, these findings suggest that SCR survey design should focus less on avoiding bias and more on achieving adequate precision.

Subsequent work shifted attention from assessing how SCR performs under alternative designs to explicitly constructing designs optimised for different criteria. Efford and Boulanger (2019) derived a fast, deterministic approximation to the coefficient of variation of estimated density, showing that precision is closely related to the minimum of expected number of detected individuals *E*[*n*] and the expected number of recaptures 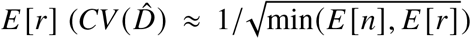. This result made it possible to rapidly evaluate and compare candidate designs without needing computationally intensive simulation. Building on this idea, Durbach et al. (2021) and Dupont et al. (2021) used genetic algorithms to optimise different criteria linked to SCR data to produce proposed designs. These approaches demonstrated that non-regular designs can match or outperform traditional grids in terms of precision, and algorithmically generated designs may offer an advantage in complex landscapes where systematic placement of detectors is difficult or infeasible.

However nearly all previous design work is based on a single “focal” set of SCR parameters. In practice, however, many surveys aim to monitor populations comprising distinct groups with marked differences in detection parameters, such as different sexes, age classes, or species. For example, male leopards typically range over areas several times larger than females resulting in very different sex-specific detection kernels. Similarly, multi-species camera-trap surveys often target species that differ markedly in body size and movement behaviour and designs optimised for one species may perform poorly for another (Foster and Harmsen 2012), yet there is currently little guidance on how detector layouts should be chosen when inference is required for multiple targets simultaneously. For convenience we henceforth use the term “group” to refer to any kind of subgroup (e.g. species, sex) that might be surveyed with a single array.

Clustered designs have been suggested as a practical way of accommodating heterogeneity in movement by combining wide inter-cluster spacing, suitable for wide-ranging individuals, with smaller within-cluster spacing that improves recaptures for animals with more restricted movement (Clark 2019), but without empirical testing. More recently, a two-stage approach has been proposed for the design of multi-species SCR surveys, whereby an initial subset of detectors is placed to maximize the number of individuals detected from the wider-ranging group, and the remaining detectors are then positioned to maximize spatial recaptures of the other group (Curveira-Santos et al. 2026). However, its performance was only assessed relative to regular grid designs, which are known to be suboptimal for species with different movement characteristics, leaving its comparative performance against more competitive design strategies unclear. Thus, the identification and systematic comparison of alternative SCR design strategies that explicitly account for multiple species with differing movement and detection parameters remain limited. In this paper, we address this gap through a simulation study of SCR design that considers scenarios with two types of animals that differ in movement scale and detection, motivated by sex-specific inference in large carnivores and by multi-species monitoring surveys. We evaluate the performance of (i) systematic grid designs including target-specific optimal spacing; (ii) clustered designs informed by contrasting movement scales; (iii) recently proposed two-stage designs; (iv) genetic algorithm–based designs optimising both the criteria of Durbach et al. 2021 and Dupont et al. 2021; and (v) lacework designs. By comparing bias, precision and coverage for both groups across the various designs, we aim to identify design principles that remain effective when SCR studies target more than one set of ecological parameters, and to provide practical recommendations for practitioners.

## 2 Methods

### 2.1 Notation

Spatial capture-recapture methods combine a spatial model that quantifies animal activity center density at all points in the survey region S with an encounter model that quantifies the expected detection frequency or detection probability, given the activity center and detector locations. Activity centers are assumed to be generated by a Poisson point process with intensity at point **s** given by *D*(**s**). We assume that detections are made at *K* proximity detectors that do not interfere with detected individuals e.g. camera traps, so that data can be collapsed into counts over a single occasion (Efford 2025). The hazard of detection for an animal with activity center at **s** is given by the hazard half-normal encounter rate function 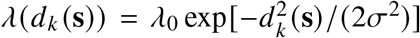, where *d*_*k*_ (**s**) is the distance between detector *k* = 1, …, *K* and activity center location **s**, *λ*_0_ is the expected encounter rate at a detector located at the animal’s activity center (*d*_*k*_ (**s**) = 0), and *σ* is a spatial scaling parameter controlling how quickly the encounter rate function decreases with distance.

For count detectors the expected number of detections of a single animal with activity center in S is given by 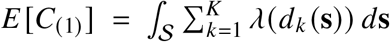 and the total number of detections collected by the survey given by 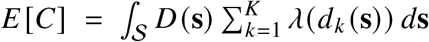 (Efford and Boulanger 2019). If density is constant over S then *E* [*C*] = *D* × *A* × *E* [*C*_(1)_] where *D* ≡ *D*(**s**) and *A* is the area of S. The expected number of individuals detected at least once is given by 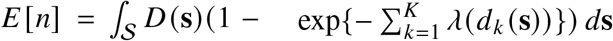 (Efford and Boulanger 2019). The expected number of recaptures is then simply *E* [*r*] = *E* [*C*] − *E* [*n*]. When referring to quantities for a specific group *g* = 1, …, *G* we use the superscript (*g*) i.e. *D*^(*g*)^, 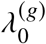, *σ*^(*g*)^. For convenience, we assume that groups are ordered by ascending values of *σ*^(*g*)^ i.e. *σ*^(1)^ ≤ *σ*^(2)^ ≤ …, ≤ *σ*^(*G*)^.

### 2.2 Design approaches

#### 2.2.1 Grid Designs

Regular grid designs are commonly used in SCR surveys and refer to detector layouts that follow a regular pattern with constant spacing *h* between detectors. Conventional guidelines are to space detectors *κσ* apart, with 1 < *κ* < 2 (Efford 2025) but it is also possible to use numerical methods to choose the spacing that maximizes the approximate precision of density by balancing the number of detected individuals with the number of recaptures (Efford and Boulanger 2019).

These grid designs pose obvious challenges for surveying multiple groups where each group has its own *σ*^(*g*)^, and these may vary substantially between groups. Using extrema *h* = *κ* max_*g*_ *σ*^(*g*)^ and *h* = *κ* min_*g*_ *σ*^(*g*)^ represent obviously poor choices and effective worst cases. More reasonable options are (a) to use the arithmetic mean *h* = *κ* (1/ *G*) *σ*^(*g*)^; (b) to use the geometric mean *h* = *κ* [ П_*g*_ *σ*^(*g*)^]^1/*G*^, which for two groups gives an equal multiplicative compromise between the group-specific spacings *κσ*^(1)^ and *κσ*^(2)^, since *h*/(*κσ*^(1)^) = *κσ*^(2)^/*h*; (c) to extend optimal spacing to more than one group by choosing the spacing that optimizes some function aggregating the group-specific precisions 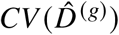. Two candidates are to maximize the mean precision over groups, giving spacing 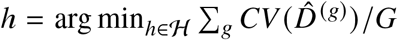, or to maximize the worst precision over groups, which gives 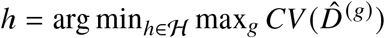.

#### 2.2.2 Cluster Designs

Cluster designs typically systematically place small detector arrays e.g. 2 × 2 or 3 × 3 regular grids, with small spacing between detectors on a grid with relatively large spacings between clusters. With their variable spacing, these designs hold obvious appeal for surveying heterogeneous populations, especially when there are two groups. For a fixed number of detectors, the main design decisions are the choices of within-cluster and between-cluster spacings, and the number of detectors per cluster (equivalently, the number of clusters).

For two groups, it seems reasonable to choose within-cluster spacing *h*_*w*_ based on the smaller *σ*^(1)^ and between-cluster spacing *h*_*B*_ based on the larger *σ*^(2)^. Possible options are (a) to use within-cluster spacing *h*_*w*_ = *κσ*^(1)^ and between-cluster spacing *h*_*B*_ = *κσ*^(2)^, with 1 < *κ* < 2; (b) to calculate the optimal detector spacing for each group independently of the other, using the smaller of these for the within-cluster spacings and the larger for the between-cluster spacing. The choice of how many detectors to use per cluster depends largely on the number of detectors available. For a single group, the main reason to use clusters is to increase the number of detected individuals at the expense of some recaptures, and thus improve precision. Because individuals are typically not detected across different clusters, each cluster should cover at least one home range, and the distance between clusters has relatively little effect on density (Clark 2019). The situation when surveying multiple groups is different. Recaptures across clusters provide important information on the larger *σ*^(2)^, so between-cluster spacing is an important choice and there will need to be an adequate number of clusters to cover the more wide-ranging group’s homerange. Thus in general we would expect cluster designs for multiple groups to contain more clusters, each with relatively few detectors, than typical cluster designs for single populations.

It is still important that detectors within a cluster cover the smaller of the home ranges. If relatively few detectors are available then it may not be possible to do this with a constant within-cluster spacing and allow enough clusters to cover the larger of the home ranges. In this case dif-ferent clusters may use different within-cluster spacing to ensure that spatial recaptures are obtained for the more restricted species over a range of distances, and detectors within some clusters are sufficiently spaced to cover a home range. Hollow (or ring) within-cluster arrangements offer another alternative.

Cluster designs are less well-suited to surveying more than two groups, at least in their standard form. A reasonable approach may be to choose the within- and between-cluster spacing based on the smallest and largest values of *σ*^(*g*)^, and then check that expected precision for intermediate groups is acceptable via simulation.

#### 2.2.3 Lacework Designs

Lacework designs consist of two sets of equally-spaced linear detector arrangements that cross at right angles to each other to form a lattice (Efford 2025). Designs are specified by two spacing parameters – the distance between detectors within a line, and the distance between parallel lines– and a third distance parameter that drops any detectors in the lattice that are more than that distance value away from where two lines intersect, which reduces the number of detectors needed. In such cases these designs produce cluster-like designs, with detectors within clusters arranged in a cross pattern rather than a grid.

Lacework designs have not been applied in practice but hold appeal for surveying multiple groups, especially with two groups, for the same reasons that cluster designs are attractive. While there are no mechanisms or guidelines for choosing within-line and between-line spacings, the arrangement of detectors between and within lines is closely related to cluster designs and so the same selection strategies described in the previous section can be used for lacework designs, with the third distance parameter used to constrain the number of detectors used.

#### 2.2.4 Algorithmic Designs

Optimal designs are those that optimize an established statistical criterion. The optimal design of SCR surveys has been infeasible because optimality criteria must be computed by repeatedly fit-ting SCR models to many simulated capture histories, each requiring numerical maximization of the likelihood. Detector layouts are usually found by iterative numerical search, requiring thousands of criterion evaluations, making the problem computationally prohibitive. Some design approaches have avoided this problem by replacing standard optimality criteria with faster but ad hoc quantities (“`En2`” designs maximizing the number of animals detected on at least two detectors (Dupont et al. 2021)) or optimizing occasionally unreliable approximations of statistical criteria (“`min(n,r)`” designs minimizing approximate 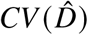, (Durbach et al. 2021)). These quantities can be efficiently calculated given a detector layout and encounter rate function, requiring initial guesstimates for *λ*_0_ and *σ*, and designs are conditional on these values. Density must also be specified although, for a single population, `En2` and `min(n,r)` designs are insensitive to this choice, which scales all derived quantities by a constant factor. Following Efford (2025), we call these designs algorithmic rather than optimal.

Simulations have shown that, for single groups, algorithmic designs offer performance that is broadly comparable to systematic designs (Dupont et al. 2021; Durbach et al. 2021). Their flexibility can be a benefit when surveying irregular survey regions not amenable to systematic layouts, but they typically produce designs that are more clustered, and less spatially balanced, than systematic designs, which will inflate variance when there is unmodelled spatial heterogeneity in density or detection (Efford 2025, Chapter 8). The same benefits and drawbacks apply when surveying multiple groups.

As with other designs, extending algorithmic designs to more than one group requires some choice of how to aggregate over groups – here, how to formulate a single objective function for all groups. Optimizing performance for one of the groups, like for regular grids, is an obviously poor alternative useful only as a baseline for comparison. Better options are (a) to base designs on the mean 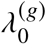 and *σ*^(*g*)^ over groups, (b) extend the algorithmic approach to multiple groups by optimizing a function that aggregates the group-specific objective function values, denoted *z*^(*g*)^, for example the mean _*g*_ *z*^(*g*)^/*G* or worst performance over groups (min_*g*_ *z*^(*g*)^ for `En2` designs; max_*g*_ *z*^(*g*)^ for `min(n,r)` designs).

#### 2.2.5 Two-Stage Designs

Curveira-Santos et al. (2026) recently proposed an algorithmic design for surveying multiple species. Their approach divides the set of available detectors into two subsets, whose placement is decided sequentially. In the first stage, the first batch of detectors is placed to maximize the expected number of detected individuals of whichever species is most mobile (i.e. has the largest expected *σ*), nominally the “focal” species. The method implements this step by algorithmic maximization of the expected number of detected inviduals, i.e. *E* [*n*], although systematic or space-filling designs would achieve the same purpose. In the second stage the detector locations from the first stage are considered fixed, and the remaining detectors are placed to maximize the expected number of spatial recaptures obtained across the remaining species.

### 2.3 Simulation study

#### 2.3.1 Overview

For each of 500 simulation replicates, detector configurations were generated using each of the proposed design approaches, with either 40 or 120 detectors. Then for each design configuration activity centre locations were simulated for an animal population consisting of two different groups, using group-specific density and detectability parameters (S2.3.2) over a defined survey region (S2.3.3). Lastly, we simulated a capture history for the simulated population and fitted maximum likelihood SCR models to each capture history to estimate group-specific parameters (S2.3.5). We used estimated parameters over all 500 replicates to assess the relative bias, precision, and coverage of estimators obtained from each design approach (S2.3.6). We discarded replicates that suggested estimation problems arising from pathologically poor designs, usually a total or near-total absence of spatial recaptures, recording the proportion of discarded replicates for each design. Specifically, the replicate for a particular group was discarded if any of the parameter estimates or standard errors were infinite or unavailable, if the parameter estimate was more than ten fold away from the true value, or if the standard error was more than 100 times its corresponding parameter estimate.

#### 2.3.2 Group-specific density and detectability

Activity center locations and detection data were simulated for two groups with very different parameter values *D*^(*g*)^, 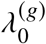 and *σ*^(*g*)^. Estimator precision is strongly dependent on the number of individual animals and spatial recaptures obtained by a survey, but calculating expectations of these depend on detector layout (Efford and Boulanger 2019). To encourage similar expected precision in each group before knowing the detector layout, we selected parameters so that 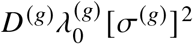 was equal for both groups, on the basis that for *K* count detectors with half-normal hazard, the integrated hazard over a sufficiently buffered state space is approximately 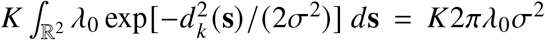, and therefore that under constant density the expected number of detections is proportional to *Dλ*_0_*σ*^2^. Note that this differs from the realized expectation *E* [*C*] that can be calculated once the detector configuration is chosen.

We simulated one group to be relatively much more wide-ranging than the other (*σ*^(1)^ = 200,*σ*^(2)^ = *Rσ*^(1)^), with *R* = 15. We used a compensatory *R*-fold difference in the baseline encounter rate 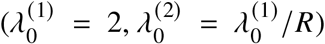, and density 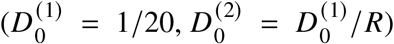. With *R* = 15, 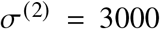, 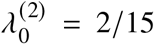, and 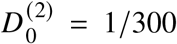. While not entirely unrealistic if one considers the two groups to be different species, we chose this relatively extreme scenario to place an upper bound on the differences between design approaches, particularly between those that account for multiple groups and those that do not. To assess sensitivity of grid designs with a single spacing to differences between groups, we assessed the performance of these designs for *R* = 1, 2, …, 15.

#### 2.3.3 Survey area

Parameter estimation by maximum likelihood SCR methods requires numerical integration of the likelihood over a fine mesh of points called the habitat mask or state space, which here also defines the study region. We used a 225 × 225 habitat mask with spacing 200m (the smaller *σ*^(1)^) and buffer 9km (three times the larger *σ*^(2)^ of 3km). This defines a 45 × 45km study area, of which the interior 27 × 27km area is available for detector placement.

#### 2.3.4 Generating detector layouts

Within each simulation replicate, we generated proposed designs with either 40 or 120 detectors. The choice of 40 detectors was motivated by many published SCR camera-trap studies having used a similar number of detectors (Tobler and Powell 2013), whereas 120 detectors is a threefold increase representing intensive sampling. Figure 1 shows one realization of the simulated designs for each of the approaches, using 40 detectors (only one grid design is shown), and Figure 2 shows the samefor 120 detectors.

**Figure 1:**
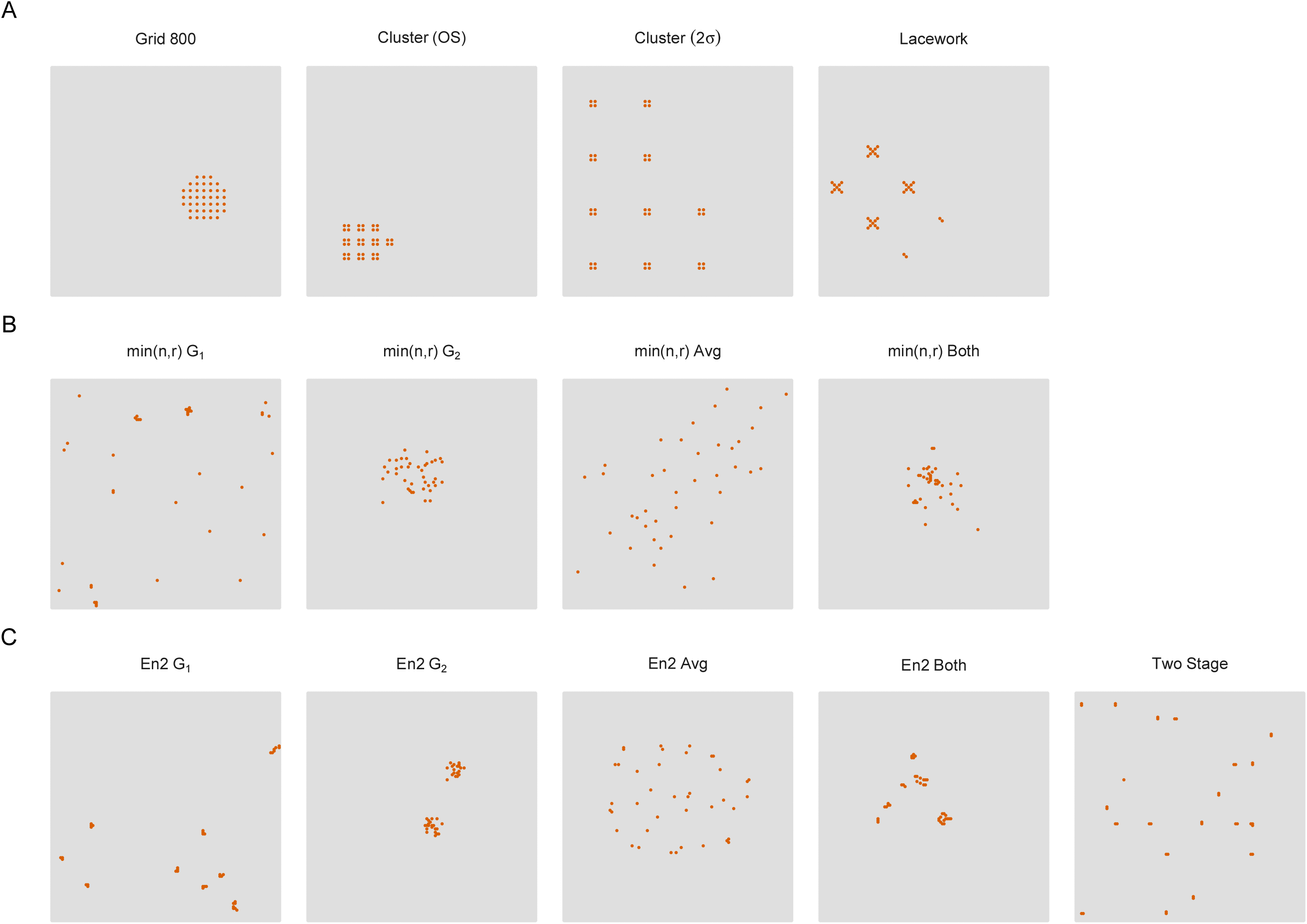
A realisation of each design used in this work for 40 traps. Panel A only includes one of the grid designs and so shows four systematic designs, panel B shows the designs optimising min(n,r), and panel C the designs optimising En2 and the two-stage design. The plotted area excludes the buffer of 9 km.

**Figure 2:**
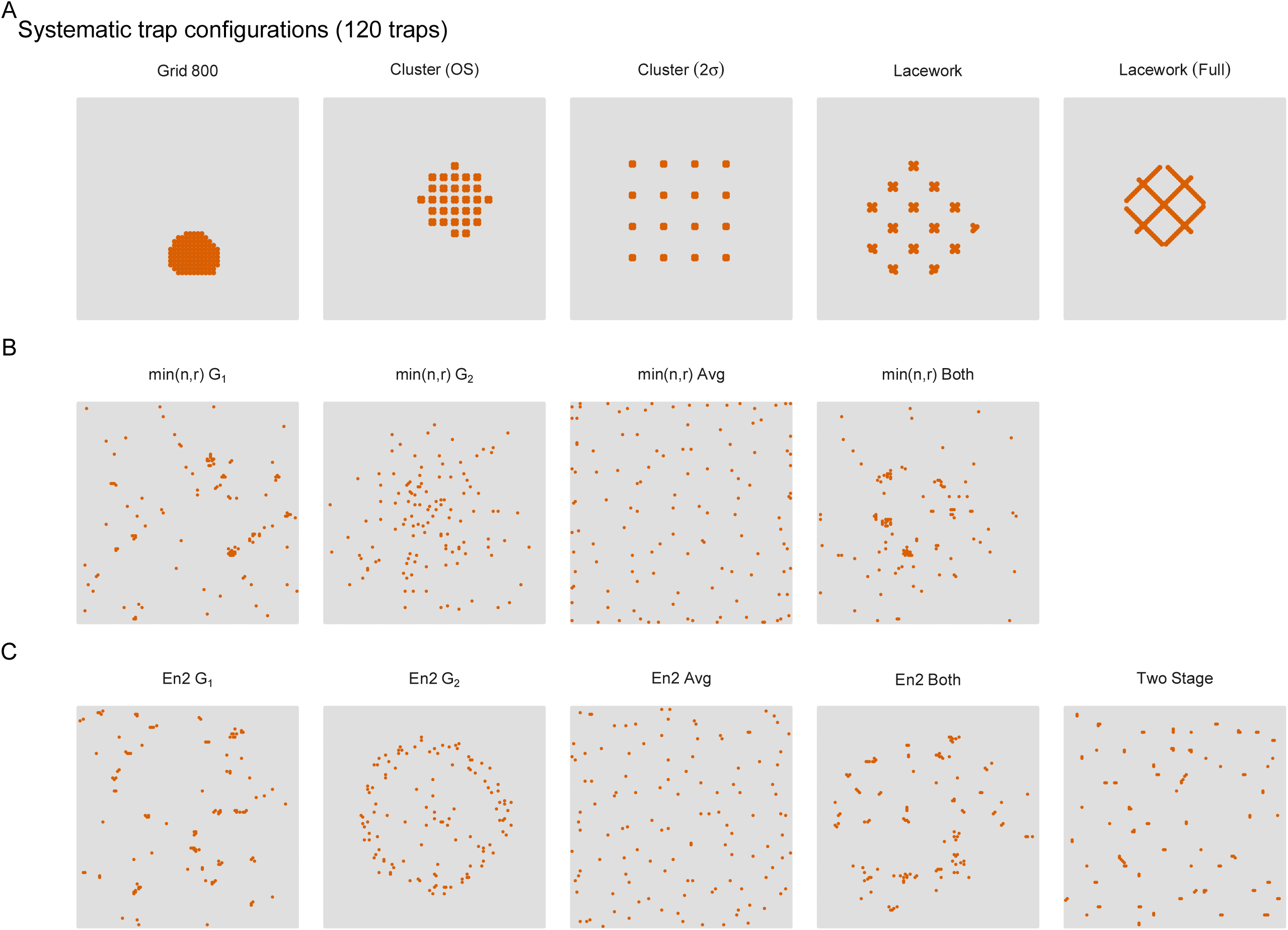
A realisation of all the designs used in this work with 120 traps. Panel A only includes one of the grid designs but includes two lacework designs and so shows five systematic designs, panel B shows the designs optimising min(n,r), and panel C the designs optimising En2 and the two-stage design. The plotted area excludes the buffer of 9 km. Note that the 2 *σ* cluster approach was only able to place 64 detectors in the allowable space.

Regular grid designs were generated using a spacing of either 800 m, the approximate geometric mean of the two *σ*s (being 4 × *σ*^(1)^ and just over 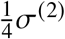) – or 600 m for 40 traps and 1,300 m for 120 traps, obtained from numerical minimization of max 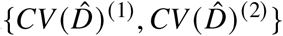.

Cluster designs used two spacings. One used 2*σ*^(1)^ for the within-cluster spacing and 2*σ*^(2)^ for between-cluster spacing, while the other calculated the optimal detector spacing for each group independently of the other, and used the smaller of these for the within-cluster spacings (500m for 40 traps and 600m for 120 traps) and the larger for the between-cluster spacing (1,000m or 1,700m). To obtain enough clusters, and given the high *λ*_0_ value of the group with more restricted movement, we used small 2 × 2 clusters. With 40 and 120 detectors available this led to 10 or 30 clusters respectively. Clusters were placed in as close to a square array as possible.

We simulated two lacework designs with the same spacings as the cluster designs i.e. 2*σ* spacings 400m and 6000m, and “optimized” spacings 500m / 1,000m for 40 traps and 600m / 1,700m for 120 traps. Given these spacings, we used trial and error to choose a value for the 3rd parameter (called `radius`) that resulted in at least several clusters being formed. We ran a third lacework design for 120 traps that did not use the radius argument, as this resulted in a lattice pattern that spanned an area of approximately the size of a home range for the wider ranging species (see top right panel of Figure 2).

Algorithmic designs were simulated by optimizing either the `min(n,r)` or `En2` criteria using ge-netic alorithms. Optimization was implemented with the `GAoptim()` function in the `secrdesign` package (Efford 2026b). Both algorithms were optimised for different parameter choices: (a) optimizing the design for the set of parameter values from group 1 only, (b) optimizing the design for group 2 only, (c) optimizing the design for the average values across the two groups, (d) optimizing the design for the average of objective function values across the two groups, or (e) optimizing the design for the worst-performing objective function values across the two groups. Both approaches performed very similarly and for the sake of brevity we only report performance from (d).

Lastly, we applied the recently developed two-stage method of Curveira-Santos et al. (2026). This approach splits the available number of detectors and in the first stage uses a space filling method to place the first set of traps. In the second stage the detector locations from stage 1 are considered fixed and the remaining traps are placed by conducting an optimisation. We allocated an equal number of detectors to each stage, using the Generalized Random Tessellation Stratified (GRTS) algorithm (Stevens and Olsen 2004) to produce a space-filling design for stage 1, and optimised `En2` with the genetic algorithm for stage 2.

#### 2.3.5 Capture histories and model fitting

For each of the 500 replicates and for each design approach, we simulated activity centers and capture histories for both groups using the parameters above. Maximum likelihood SCR models were then used to estimate group-specific parameters.

#### 2.3.6 Comparing designs

Based on 500 replications, for each parameter 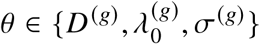 we calculated (a) the relative bias, 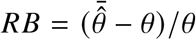, where 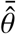 is the empirical mean of the parameter estimate 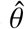 and *θ* is the true parameter value, (b) precision,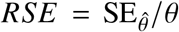, where 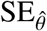 is the empirical standard deviation of parameter estimate 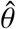, and (c) coverage, calculated as the proportion of replicates in which *θ* fell in the interval 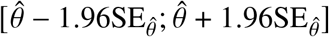

#### 2.3.7 Software

Simulations were implemented in R v4.5.2 using the secr (Efford 2026a) and secrdesign (Efford 2026b) packages, with some customized extensions used to accommodate two groups. Functions within these packages were used to generate grid designs (`make.grid()`), lacework designs (`make.lacework()`), and algorithmic designs (`GAoptim()`). Cluster designs were generated by a small extension to `make.grid()`. Two-stage designs used the `grts()` function from the spsurvey package for the space-filling first stage, and a custom extension of `GAoptim()`) allowing some detectors to be held fixed for the second stage. The numerical minimization of max 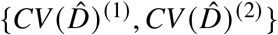 used to choose an optimized regular grid spacing was done using a search over values generated with the `minnrRSE()` function in secrdesign. Algorithmic designs optimizing the sum of objective function values over the two groups were generated by extending `GAoptim()` to allow for two sets of input parameters to be specified, and for expected sample sizes to calculated for each group. Activity centers and capture histories were simulated using `sim.popn()` and `sim.capthist()` respectively, and models were fitted using a “group” model in `secr` (i.e. `secr.fit()` with the groups argument specified). Computations were performed using the University of Cape Town ICTS High Performance Computing infrastructure (UCT HPC 2023). All code and output are available at https://github.com/gdistiller/Designing-SCR-surveys-for-multiple-groups.

## 3 Results

The results of the simulations are reported for density in Figure 3 and for *σ* in Figure 4. A similar figure for *λ*_0_ can be found in the supplementary material. There were relatively few cases with 40 traps where replicates were discarded with these proportions ranging from 0.2% for group 2 (with En2-G1) to 7.6% for group 2 (with Grid (2*σ*)), further details are in the supplementary material. All replications with 120 traps were retained apart from the `Grid (OS)` design that used both sets of parameters to determine the optimal spacing (of 1,300m). That case resulted in 39% of group 1’s estimates being discarded, and poor performance on several of the metrics for the remaining replicates.

**Figure 3:**
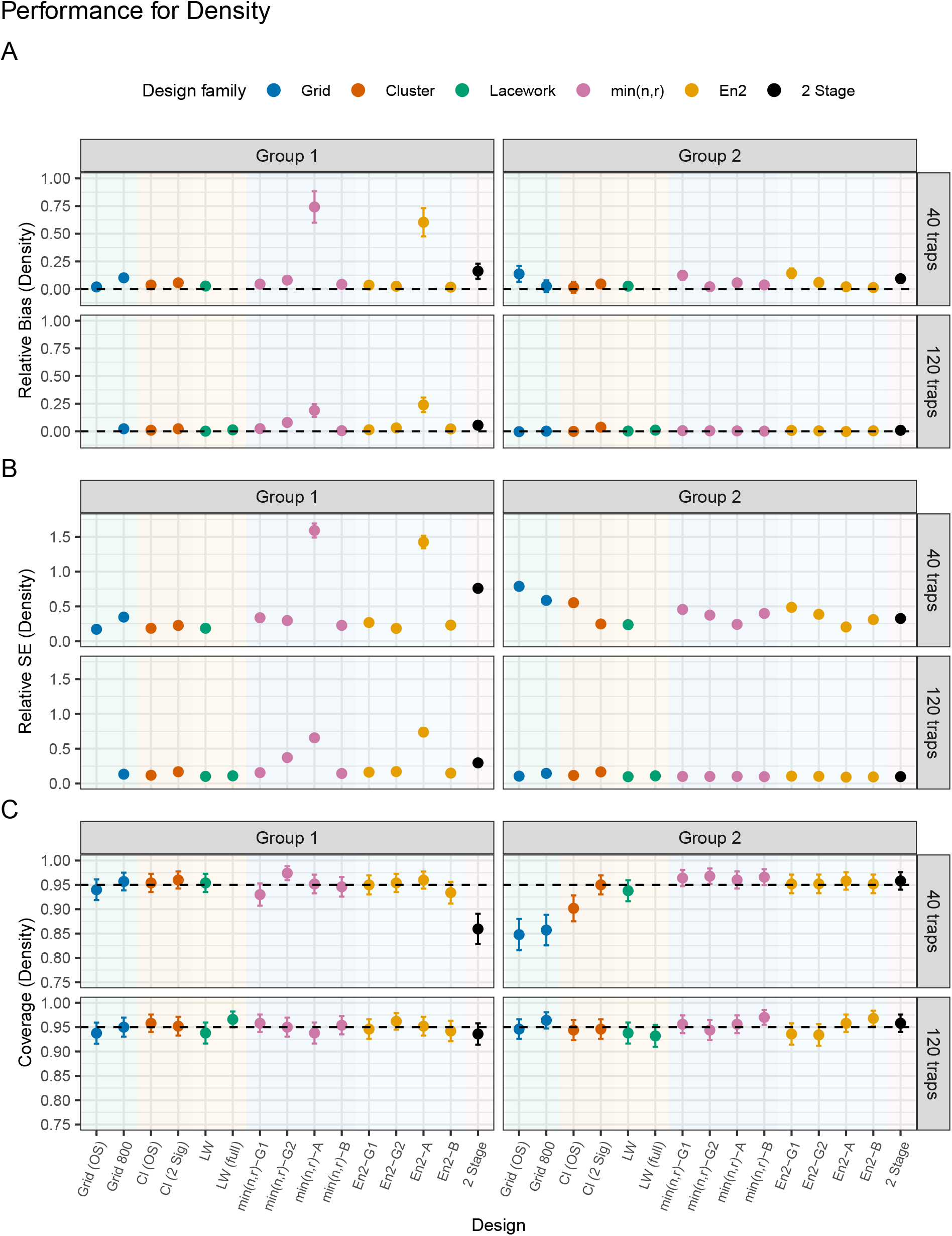
Relative bias (A), relative standard error (B), and coverage (C) of 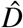 for each design. OS = optimal spacing, G1/G2 = group-specific parameters; A = optimizing average; B = minmax or maxmin optimization. Missing points indicate either extremely poor performance (`Grid (OS)`, 120 detectors) or infeasible designs (`LW (full)`, 40 detectors). Vertical bars denote two standard errors.

**Figure 4:**
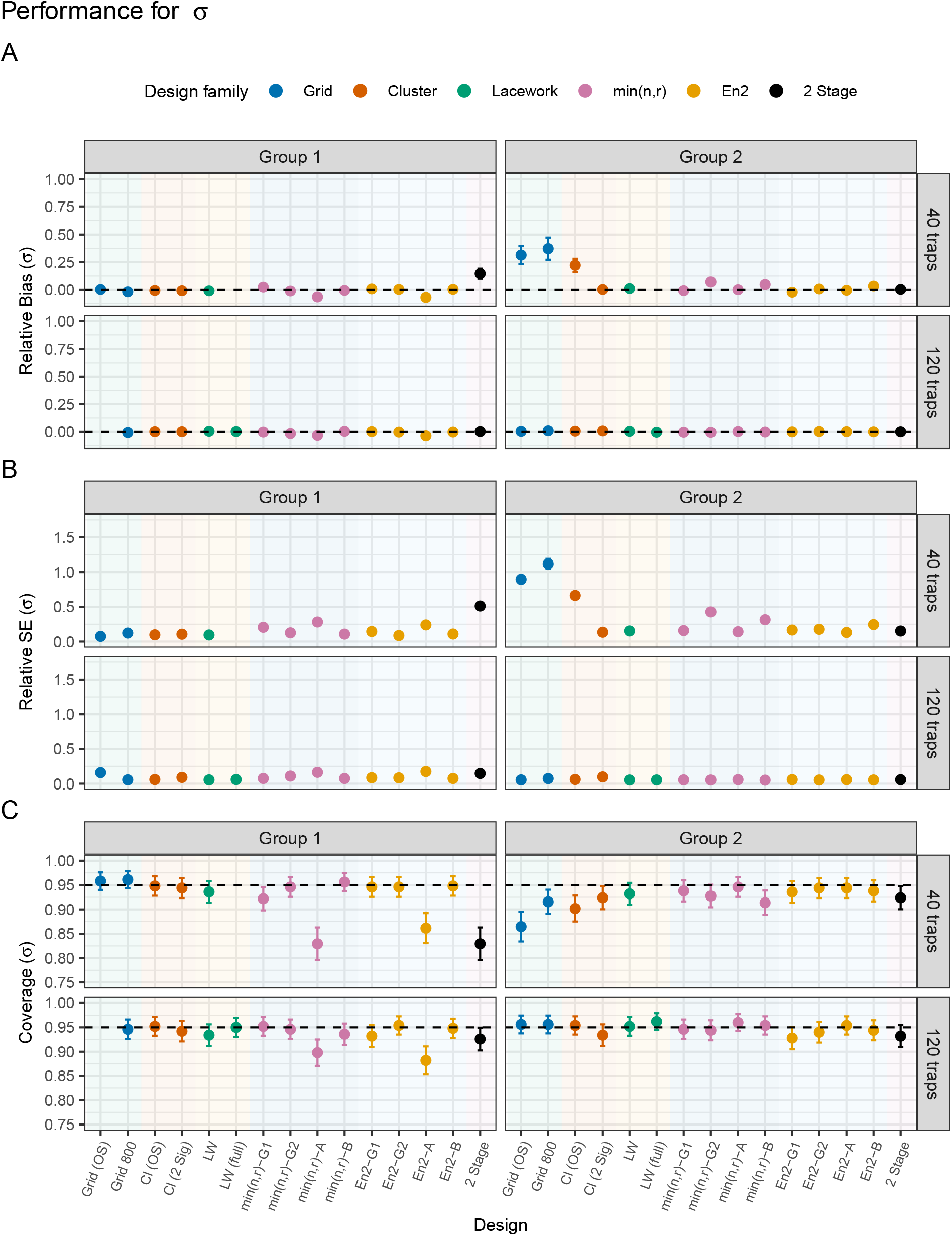
Relative bias (A), relative standard error (B), and coverage (C) of 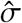 for each design. OS = optimal spacing, G1/G2 = group-specific parameters; A = optimizing average; B = minmax or maxmin optimization. Missing points indicate either extremely poor performance (`Grid (OS)`, 120 detectors) or infeasible designs (`LW (full)`, 40 detectors). Vertical bars denote two standard errors

### 3.1 Bias

When using 40 traps, the relative bias in density was low across all the different design approaches, apart from when using the `min(n,r)` or `En2` approach with parameter values that were averaged across the two groups which led to positive bias for both groups, though the bias was substantially worse for group 1. There was also slight positive bias for group 2 when using an optimal spacing grid and for both groups when using the two-stage design.

The *σ* parameter was also generally estimated with little bias. However using a grid design led to positive bias of around 40% for group 2, and using a cluster design with optimal spacings to a positive bias of just under 25% for that group.

The estimator for *λ*_0_ was unbiased across all scenarios. Increasing the number of traps to 120 led to slight positive bias in estimated density for group 1 when optimising with average parameter values. All other signs of bias in all parameters disappeared with the extra detectors.

### 3.2 Precision

With 40 traps, most of the approaches led to slightly larger density RSEs for group 2, though the worst precision was for group 1 when using the algorithmic approaches with average values followed by the two-stage design. For group 2 the highest RSE in D was from the grid designs where it exceeded 0.5 for both spacings, and the cluster design with optimal spacing was just behind these followed by the algorithmic approaches for most of the parameter choices. The 2*σ* cluster design, the lacework cluster design, and the algorithmic approaches with average values had the lowest RSEs for group 2.

We used an extreme difference between the groups (*R* = 15) and in such a case it is clear that a grid with consistent spacing does not work well (also see coverage results in S3.3). We also explored how the RSE for density changed with the difference between groups, when using the geometric mean approach to determine the spacing for the grid, and the results are shown in Figure 5. It is clear that RSE for group 2 increases with increased spacing between the traps. On the other hand the RSE for group 1 is fairly flat until one gets a difference of about 8 or 9 fold at which point the higher spacings associated with those *R* values resulted in a reduction in precision.

**Figure 5:**
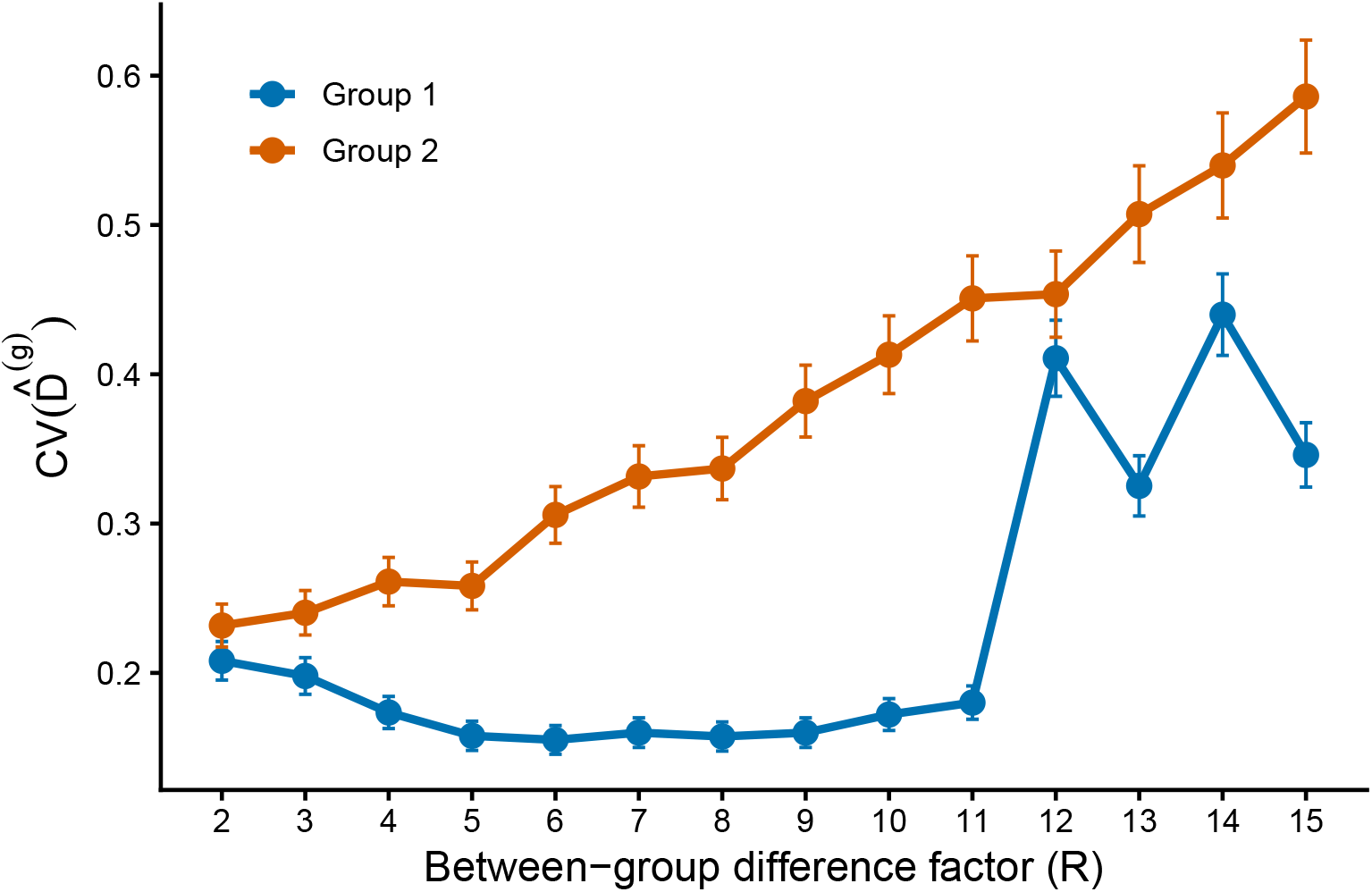
CV(*D*^*g*^) vs *R*, the factor that determines the difference between the two groups. Vertical bars represent two standard errors.

The RSE for the *λ*_0_ parameter was consistently low with little differences between the designs or the groups. For *σ* some of the algorithmic approaches and the two-stage design had slightly higher RSEs for group 1 than other designs, whereas the RSE for group 2 was highest when using the grid designs or the cluster design with optimal spacings.

As expected, precision improved across the board when increasing the number of detectors to 120, although RSE in D for group 1 was still above 0.5 when optimising with average values. RSE for *λ*_0_ and *σ* was consistently low with little differences between the groups or designs.

### 3.3 Coverage

With 40 traps, coverage in D for group 1 was good for all designs with the exception of the two-stage approach. Other than that the only approach where the interval missed the nominal level was when optimising `min(n,r)` with group 2 values, and in that case the coverage was just above the nominal level. On the other hand the coverage for D in group 2 was poor when using grids or the cluster design with optimal spacings.

For the *λ*_0_ parameter coverage was at or near the nominal level across both groups for all approaches. However for *σ* there was poor coverage (below 0.95) for group 1 when using the algorithmic designs with average values and the two-stage design. The coverage for *σ* in group 2 was also below the nominal level for the grid designs and the cluster design with optimal spacings. When using 120 traps, coverage in D and *λ*_0_ was good for all cases. However the group 1 coverage for *σ* still remained below the nominal level when optimising with average values. Coverage for *σ* was good for all options in group 2.

## 4 Discussion

Camera traps are increasingly used to obtain information on multiple species from a single field survey, and secondary analysis of non-target detections is widely encouraged because it can increase the conservation value obtained from costly sampling effort (Scotson et al. 2017; Delisle et al. 2021). However, this value is likely to be greater when multi-species use is considered during survey design rather than treated only as a post-survey opportunity. Detector layouts are tied to species-specific movement and detection processes, so that a design that performs well for one species may do poorly on another. Multi-species designs must balance competing spatial requirements, yet there is little guidance on how this should be done. Here, we evaluate practical ways of extending familiar single-species designs to accommodate two heterogeneous target species.

Our results show that multi-species SCR surveys can perform well even when focal groups differ greatly in movement scale and encounter rate. The most reliable designs were systematic layouts containing more than one characteristic spacing. Cluster and lacework designs performed consistently well, while algorithmic designs achieved similar performance only when their objective functions explicitly represented both groups. In contrast, designs based on a single compromise spacing, a single group’s parameters, or a sequential allocation of detectors were less reliable.

Differences among many designs were relatively modest, consistent with the general robustness of SCR estimators to detector placement. Provided that a design detects enough individuals, obtains sufficient spatial recaptures, and covers an area large enough relative to animal movement, density estimates are often approximately unbiased across a wide range of layouts (Sollmann et al. 2012; Tobler and Powell 2013; Sun et al. 2014; Fleming et al. 2021). Large improvements in precision are also intrinsically difficult to obtain. The approximation 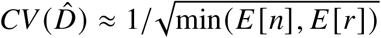 implies strong diminishing returns: halving the coefficient of variation requires a fourfold increase in whichever of the expected number of detected individuals or recaptures is limiting (Efford and Boulanger 2019). Our simulation used a 15-fold difference in movement scale specifically to create a setting in which multi-group designs should have a substantial advantage. Even then, several quite different layouts performed similarly. This suggests that for most studies there may be a broad class of effective multi-species designs, defined mainly by the need to avoid serious deficiencies for any one group.

Our results provide little support for routine use of algorithmic designs. This was a setting in which algorithmic designs might have been expected to excel, because unconstrained detector spacings provide a natural way to accommodate contrasting movement scales. The best algorithmic designs performed well but did not consistently outperform simpler systematic alternatives. Good performance required the objective function to be evaluated separately for both groups and then aggregated. Designs optimised for one group tended to perform poorly for the other, while using average values of *λ*_0_ and *σ* was less reliable and produced bias in some cases. Averaging parameter values does not preserve either the recapture requirements of the less mobile group or the spatial coverage required for the more mobile group. The `min(n,r)` and `En2` criteria produced visually different layouts but similar inferential performance. Algorithmic designs require greater computation and can produce highly clustered, spatially unbalanced layouts that are more susceptible to bias when density or detectability varies spatially (Efford 2025, Chapter 8). We therefore view them mainly as an option in irregular landscapes where systematic placement is infeasible. Where a spatially balanced systematic design can be implemented, its similar statistical performance, transparency, and robustness will make it preferable in most applications.

A regular grid with one spacing is not an adequate default when movement scales differ greatly. A spacing suitable for the less mobile group restricts the extent of the array and provides weak information on *σ* for the more mobile group, whereas a spacing suitable for the more mobile group yields few recaptures for the less mobile group. Compromise spacings avoid the worst extremes but cannot reproduce both spatial scales. This was clearest with 40 detectors, where grids gave poor precision and substantial positive bias in *σ* for the wide-ranging group. These differences largely disappeared with 120 detectors, because a larger array contains detector pairs separated by greater distances. Layout is expected to matter less when resources are abundant, but it is critical that spatial recaptures are obtained for all groups. Note that this is not guaranteed when choosing the spacing that minimizes the maximum approximate *CV* (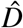) across groups, because this approximation treats spatial and non-spatial recaptures as equivalent. In our case this resulted in a pathological 120-detector design that produced extremely poor results for the less mobile group. A single spacing may be practical when differences in movement are moderate between groups. In our scenario bias and precision remained broadly acceptable for both groups until differences were greater than five-fold, although this will depend on the specific density and detection parameters of the surveyed populations, in particular whether the baseline encounter rates of the less mobile groups are sufficiently high to produce spatial recaptures at relatively large (i.e. > 2*σ*) spacings.

Cluster designs provide a natural systematic solution because within- and between-cluster spacings can be selected for different movement scales. In a two-group survey, detections within clusters provide recaptures for the less mobile group, while detections across clusters inform movement of the wider-ranging group. The main design choices are therefore the number of clusters, the number of detectors within each cluster, and the two spacings. Too few clusters may provide inadequate geographical coverage, whereas clusters that are too small or too tightly packed may provide little information about *σ*. In our simulations, small 2 × 2 clusters performed well, and separately optimising the two spacings gave no clear improvement over using simple multiples of the two *σ* values. Approximate prior knowledge of movement scales may therefore be sufficient. Lacework designs offer a related systematic alternative. Their within-line and between-line spacings can represent the smaller and larger movement scales, while crossing lines provide opportunities for recaptures at multiple distances. With only 40 detectors and an extreme difference between the two values of *σ*, however, a complete lacework lattice was infeasible – restricting the number of detectors effectively converted the lacework design into a cross-shaped cluster design. Lacework becomes more distinctive and potentially more attractive when more detectors are available or differences among movement scales are less extreme.

The two-stage design was less consistently effective. Sequential placement fixes a substantial fraction of the detector budget before all species are considered, and performance depends on the focal species, the division of detectors between stages, and the size of the survey region. A space-filling first stage only gives appropriate spacing when the number of detectors and the extent of the region happen to align with the focal species’ movement scale. The second stage also maximises the sum of recaptures over non-focal species, implicitly assigning greater weight to more abundant or detectable species. That may be appropriate in some applications, but a single-stage criterion representing all focal groups makes these trade-offs explicit and controls them more directly.

Extensions to more than two groups are straightforward for algorithmic designs, using a weighted mean or worst-case aggregation of group-specific objectives. Extensions to systematic layouts are less direct. Within- and between-unit spacings can be based on the smallest and largest values of *σ*, but performance for intermediate groups should be checked. The main risks remain insufficient recaptures for less mobile species and inadequate coverage for more mobile species. Candidate designs should therefore be evaluated using expected detections and recaptures for every focal group. As these quantities can be quickly evaluated, numerical search over potential spacing parameters is straightforward for systematic designs. As for any design, sensitivity to assumed group density and detection parameters should also be assessed.

Our simulations necessarily simplified applied multi-species surveys. We assumed homogeneous density and detection, and known group membership. Spatially varying density or encounter probability would strengthen the case for spatially balanced designs and could make the highly clustered algorithmic layouts less robust than indicated here. We also considered two groups with parameters selected so that their expected total numbers of detections were broadly comparable before detector placement. Applied surveys may instead include one common, readily detected species and another that is rare or difficult to detect, in which case overall design performance may be dictated almost entirely by the latter. We chose scenarios with camera traps in mind, and camera traps are often used to monitor species that occur at low densities and range widely (Foster and Harmsen 2012). We did not explore how the various design approaches perform when animals occur at high density and have small ranges.

In practice, we recommend identifying the smallest and largest plausible movement scales and rejecting layouts that cannot generate recaptures at the smaller scale or cover a substantial area at the larger scale. For two contrasting groups, a spatially balanced cluster or lacework design with short within-unit and wider between-unit spacing provides a reasonable default. Expected detections and recaptures should be calculated separately for each group rather than from averaged encounter parameters. Optimized spacings provide small increases in efficiency where input parameters are known relatively precisely. Algorithmic optimisation may be useful where field constraints prevent a systematic layout, but the objective should aggregate group-specific criteria directly and the resulting layout should be checked for excessive clustering.

## Notes

### Competing Interest Statement

The authors have declared no competing interest.

