## Supplementary figures for "Designing spatial capture–recapture surveys for multiple populations"

### 1 Extra Figures

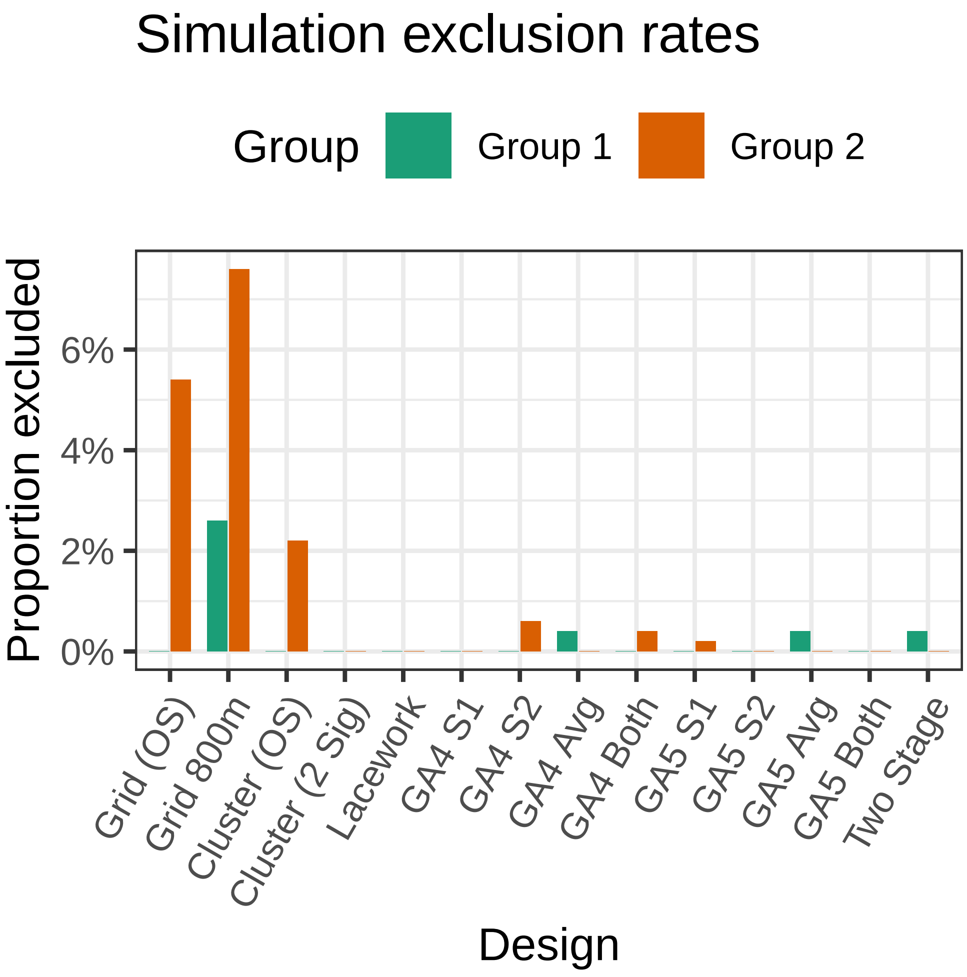


Figure 1: Proportion of the 500 replications that were discarded.

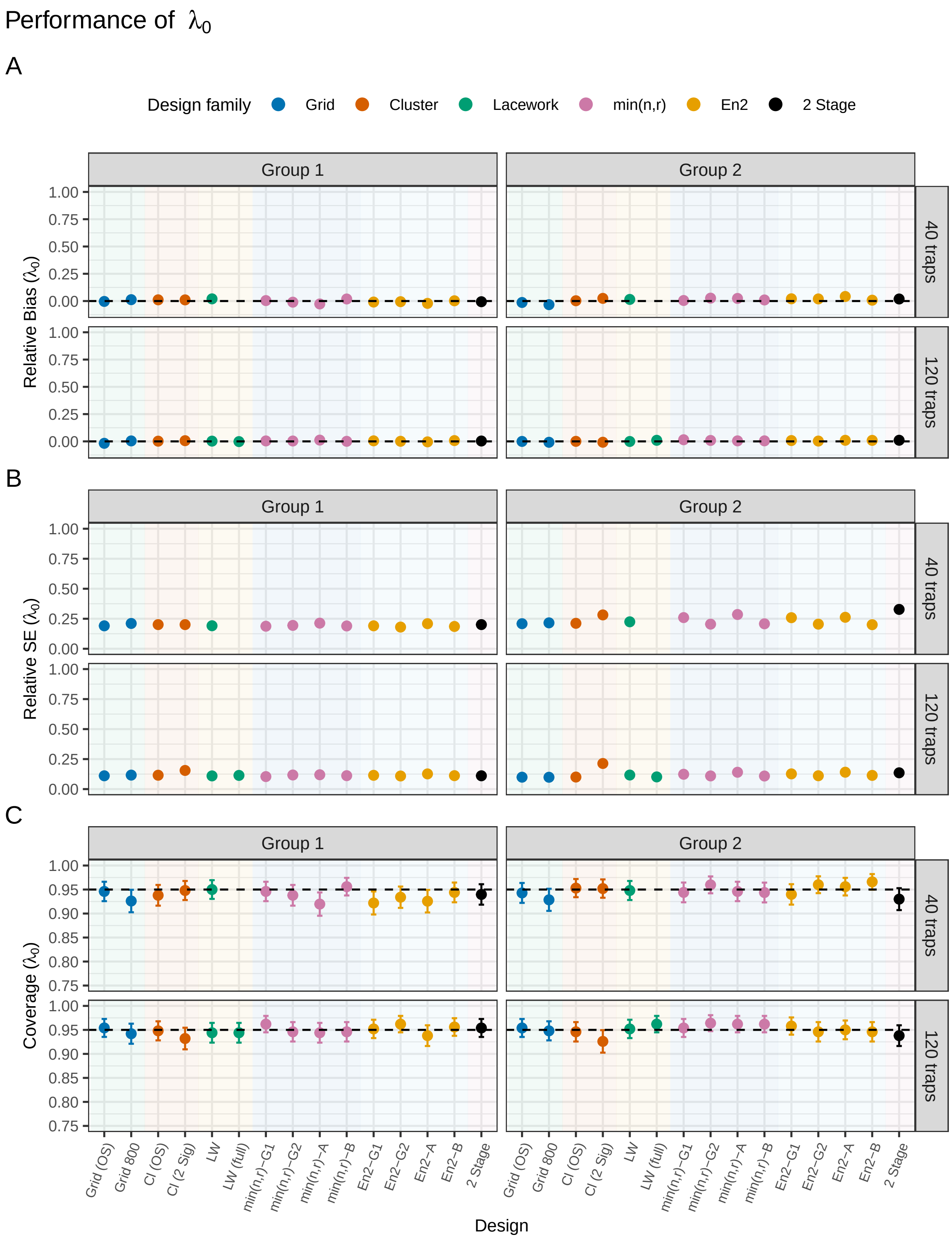


Figure 2: Performance for the $\lambda_0$ estimator for each design. Panel A shows the relative bias, panel B the relative standard error, and panel C the coverage. `Grid / Cl (OS)` refers to the grid or cluster design with optimal spacing, `LW / LW (f)` to the lacework designs with `LW (f)` indicating the lattice design that could only be applied with 120 traps, and for the algorithmic designs `G1 / G2` indicates group 1 or 2 specific parameters and `A / B` refers to using the average values or optimising with both groups. Vertical bars represent two standard errors.

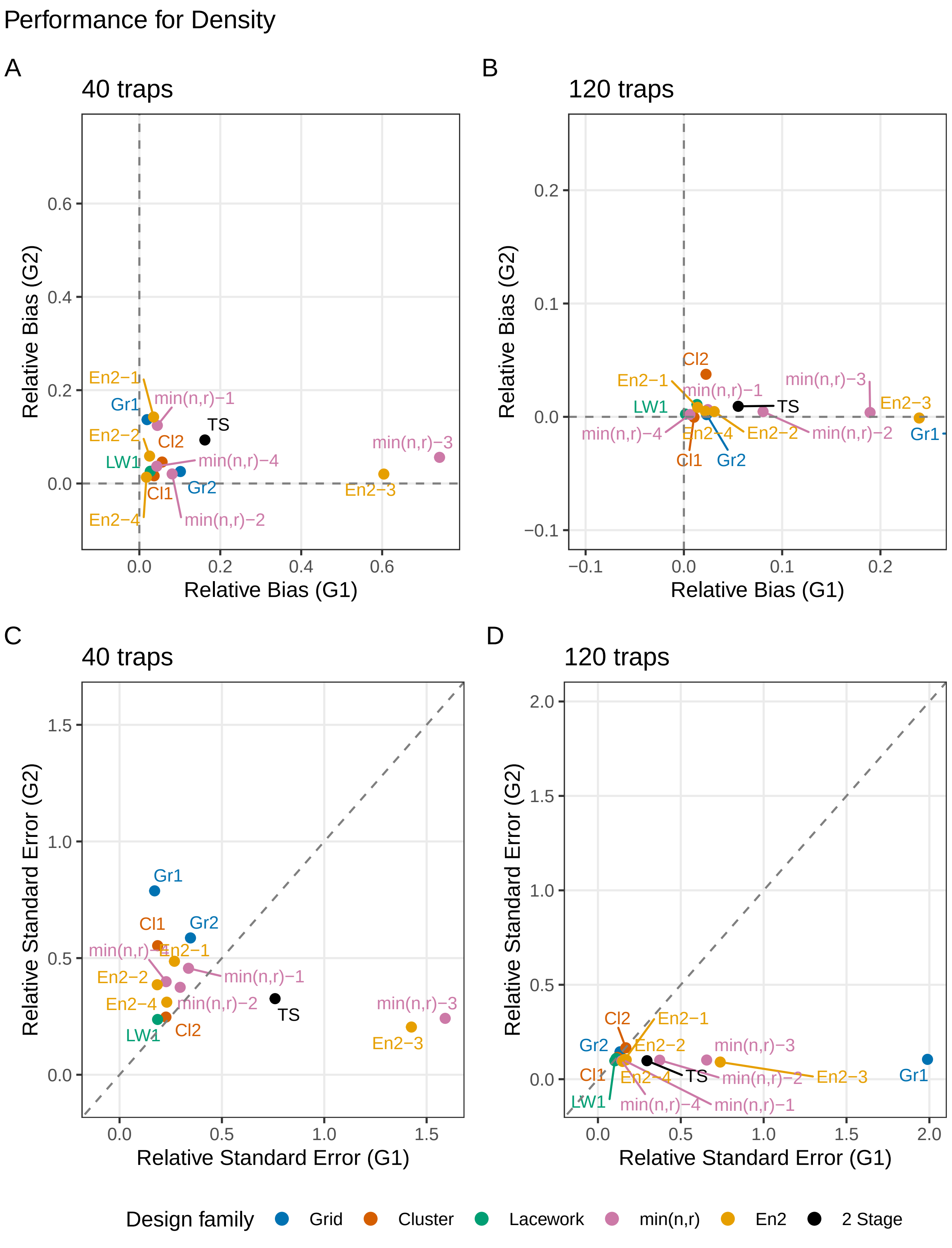


Figure 3: Two dimensional plots of RB and RSE for the density parameter. `Gr / Cl 1` refers to the grid or cluster design with optimal spacing, `LW 2` to the lacework design that could only be applied with 120 traps, and for the algorithmic designs `1/2` indicates group 1 or 2 specific parameters and `3/4` refers to using the average values or optimising with both groups.

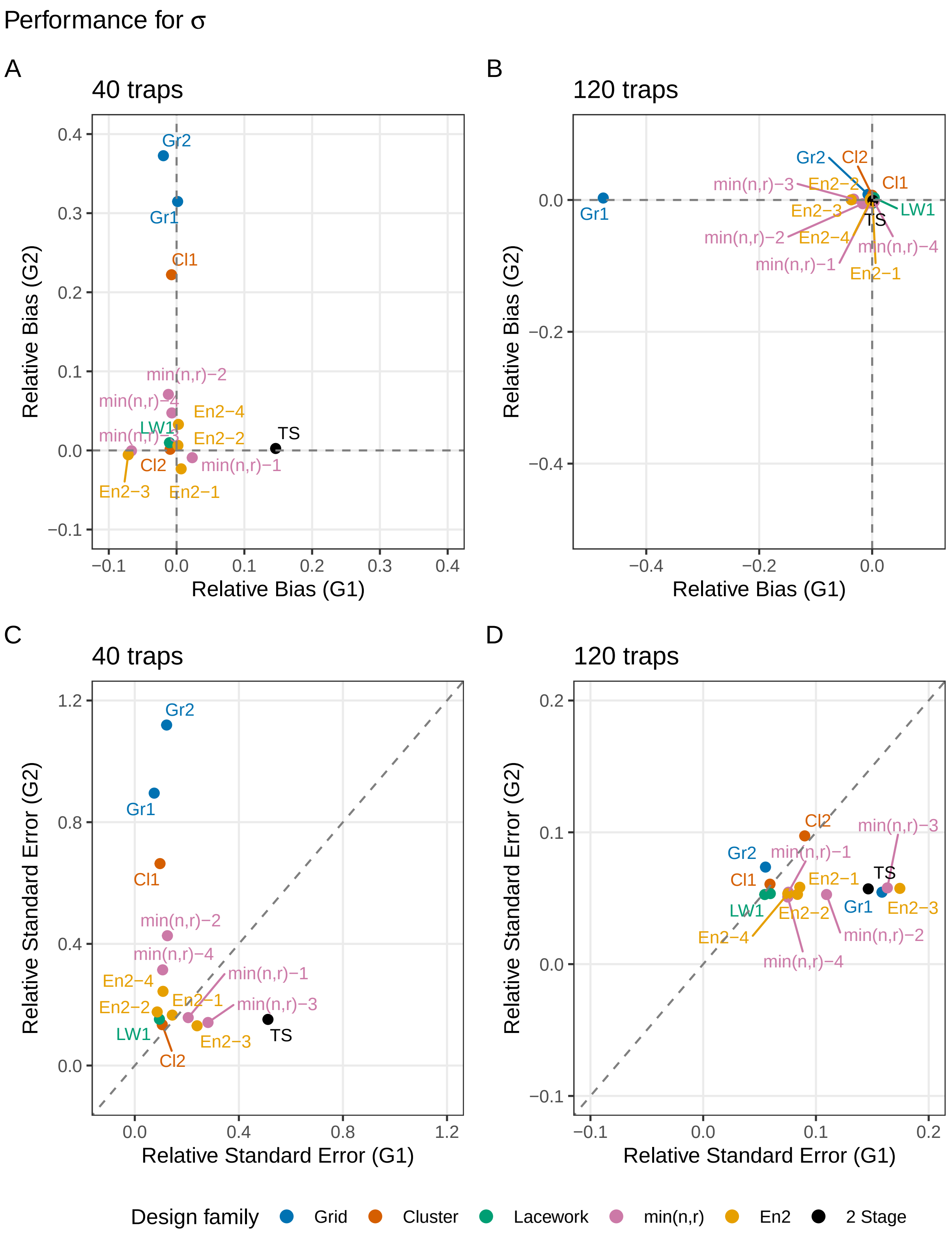


Figure 4: Two dimensional plots of RB and RSE for the $\sigma$ parameter. `Gr / Cl 1` refers to the grid or cluster design with optimal spacing, `LW 2` to the lacework design that could only be applied with 120 traps, and for the algorithmic designs `1/2` indicates group 1 or 2 specific parameters and `3/4` refers to using the average values or optimising with both groups.
